# Quantifying shape, integration, and ecology in avian pedal claws: comparing geometric morphometric and traditional metric approaches

**DOI:** 10.1101/593236

**Authors:** Brandon P. Hedrick, Samantha A. Cordero, Lindsay E. Zanno, Christopher Noto, Peter Dodson

## Abstract

Terrestrial tetrapods use their claws to interact with their environments in a plethora of ways. Birds in particular have developed a diversity of claw shapes since they are not bound to terrestrial locomotion and have heterogeneous body masses ranging several orders of magnitude. Numerous previous studies have hypothesized a connection between pedal claw shape and ecological mode in birds, but have generated conflicting results, spanning from clear ecological groupings based on claw shape to a complete overlap of ecological modes. These studies have relied on traditional morphometric arc measurements of keratinous sheaths and have variably accounted for likely confounding factors such as body mass and phylogenetic relatedness. To better address the hypothesized relationship between ecology and claw shape in birds, 580 radiographs were collected allowing visualization of the bony core and keratinous sheath shape spanning 21 avian orders. A new method was used to quantify claw shape using geometric morphometrics and was compared to results using traditional arc measurements. Neither traditional nor geometric morphometrics are capable of significantly separating bird claws into coarse ecological categories after integrating body size and phylogenetic relatedness. Further, the bony claw core and keratinous sheath are significantly integrated with one another, suggesting that they function as a single unit. Therefore, it is likely possible to compare fossil bony cores with extant keratinous sheaths after applying corrections. Finally, traditional metrics and geometric morphometric shape are significantly, yet loosely correlated, and geometric morphometric data better distinguish ecological groups in morphospace than is possible with traditional metrics. Based on these results, future workers are encouraged to use geometric morphometric approaches to study claw geometry and account for confounding factors such as body size, phylogeny, and individual variation prior to predicting ecology in fossil taxa.

## 1 Introduction

Claws are important tools that vertebrates use to interact with their environments and are used for a variety of purposes, including locomotion, clinging to surfaces, food gathering, burrowing, and in inter- and intraspecific combat. Although the relationship between claw shape and ecological mode has been examined in birds (Feduccia, 1993) and lizards (Zani, 2000; D’Amore et al., 2018), this hypothesized relationship has most often been used for predicting the ecology of extinct taxa using the claw morphology of extant taxa (Feduccia, 1993; Glen & Bennett, 2007; Fowler et al., 2009; Fowler et al., 2011; Birn-Jeffery et al., 2012; Cobb & Sellers, in review). There have been comparatively few studies focusing explicitly on extant bird claw shape, whether on the development and variability of claw morphology (Ethier et al., 2010) or the correlation between claw morphology and ecological mode (but see Pike & Maitland, 2004; Csermely & Rossi, 2006; Csermely et al., 2012). Studies that have examined extant bird taxa find conflicting levels of correlation between claw shape and ecology, with different ecological modes often having large amounts of overlap in shape (Pike & Maitland, 2004; Birn-Jeffery et al., 2012). Since birds are not bound to terrestrial locomotion, their pedal claws have different constraints than obligate terrestrial taxa and are capable of taking on a wide spectrum of shapes, such as the long recurve found in many raptorial claws. Aves also has high body size disparity ranging from the bee hummingbird (2.2 g) to the ostrich (111,000 g) (Dunning, 1993), which generates a variety of different constraints on pedal claw shape. Therefore, it would be expected that Aves would have high claw disparity likely driven by different factors in different ecological groups and that body mass would have a large impact on shape.

Previous workers examining claw morphometrics have adapted a version of the traditional morphometric arc length method first proposed by Peters and Görgner (1992) and Feduccia (1993) whereby claw shape is reduced to the angle of the claw arc. These methods have varied as to where the arc measurement was taken: on the dorsal surface of the claw (e.g., Pike & Maitland, 2004) or on the ventral surface of the claw (e.g., Feduccia, 1993), but all assume that the claw arc inscribes a circle. However, this is not the case in many bird species. Additionally, few studies have used phylogenetic comparative methods to incorporate the inter-relatedness of representative taxa in statistical analyses (Felsenstein, 1985). Finally, the majority of vertebrate claws are composed of two basic components: the distal bony ungual and the keratinous sheath that envelops the bony core. Many studies have used the shape of the bony core in extinct taxa and the shape of the keratinous sheath in extant taxa interchangeably when making functional morphological assertions (but this has long been known to be problematic–Birn-Jeffery et al., 2012). As a result, there is not yet a detailed understanding of how the bony core of the claw and the more friable keratinous sheath relate to one another and whether they can compatibly be compared.

Geometric morphometrics is a powerful technique for quantitatively analyzing shape data (Bookstein, 1991; Corti, 1993; Slice, 2007; Mitteroecker & Gunz, 2009; Zelditch et al., 2012), including recent work on claws (D’Amore et al., 2018). We develop a new method of holistically assessing claw shape using geometric morphometrics and compare the geometric morphometric approach with traditional morphometric arc measurements using radiographs of a large sample of bird pedal digit III claws from across Aves. Traditional morphometric and geometric morphometric data were extracted from both the bony core shape and keratinous sheath shape of each specimen to assess the following: (1) Is there a significant relationship between ecology and morphology in avian pedal claws after incorporating phylogeny and body size?; (2); Are the bony claw core and keratinous sheath significantly integrated with one another or do they evolve separately from one another? (3) How closely do traditional morphometric and geometric morphometric data coincide? Although the relationship between claw shape and ecology in birds has been previously examined, this is the first study to look at claw shape comprehensively using the total claw shape and not only the arc measurement of the claw, incorporating keratinous sheath shape, bony core shape, phylogenetic comparative methods, and body size.

## 2 Materials and Methods

Pedal digit III claws from 580 individuals of 145 species in 21 orders across the avian tree were x-rayed (Figure 1, S1). Digit III was used because it is the primary weight bearing toe (Glen & Bennett, 2007) and has most commonly been used in previous studies of claw shape, affording repeatability with past studies. Additional toes were not considered due to significant interdigital variation with the same foot (Fowler et al., 2009). The specimens were radiographed using a Kodex Inc. Imagex 20i with Thermo Kevex x-ray source (PXS10-16w) and Varian Digital x-ray detector. Images were taken at 40 kV and 266 μA with a spot size of 13 microns with 10.6 watts and 20 frames per radiograph. These radiographs allowed for visualization of both the bony core and keratinous sheath of each specimen. Some species were sampled at particularly high rates to assess intraspecific variation in claw shape (e.g., *Tinamus major*, n = 25). To facilitate phylogenetic comparative analyses, species means were taken for the 145 species for both traditional and geometric morphometric measures. The maximum credibility phylogeny for extant birds generated by Jetz et al. (2012) was pruned to include only the species sampled in this analysis and was used for phylogenetic comparative analyses (Figure S1). Taxa were then split into three ecological groups (predominantly predatory, predominately ground-dwelling, and flying generalists) to assess how claw shape related to ecology. To better balance sample sizes within ecological groups, flying generalists were not further split into climbing and perching birds as has been done by previous workers (Pike & Maitland, 2004; Glen & Bennett, 2007).

**FIGURE 1.**
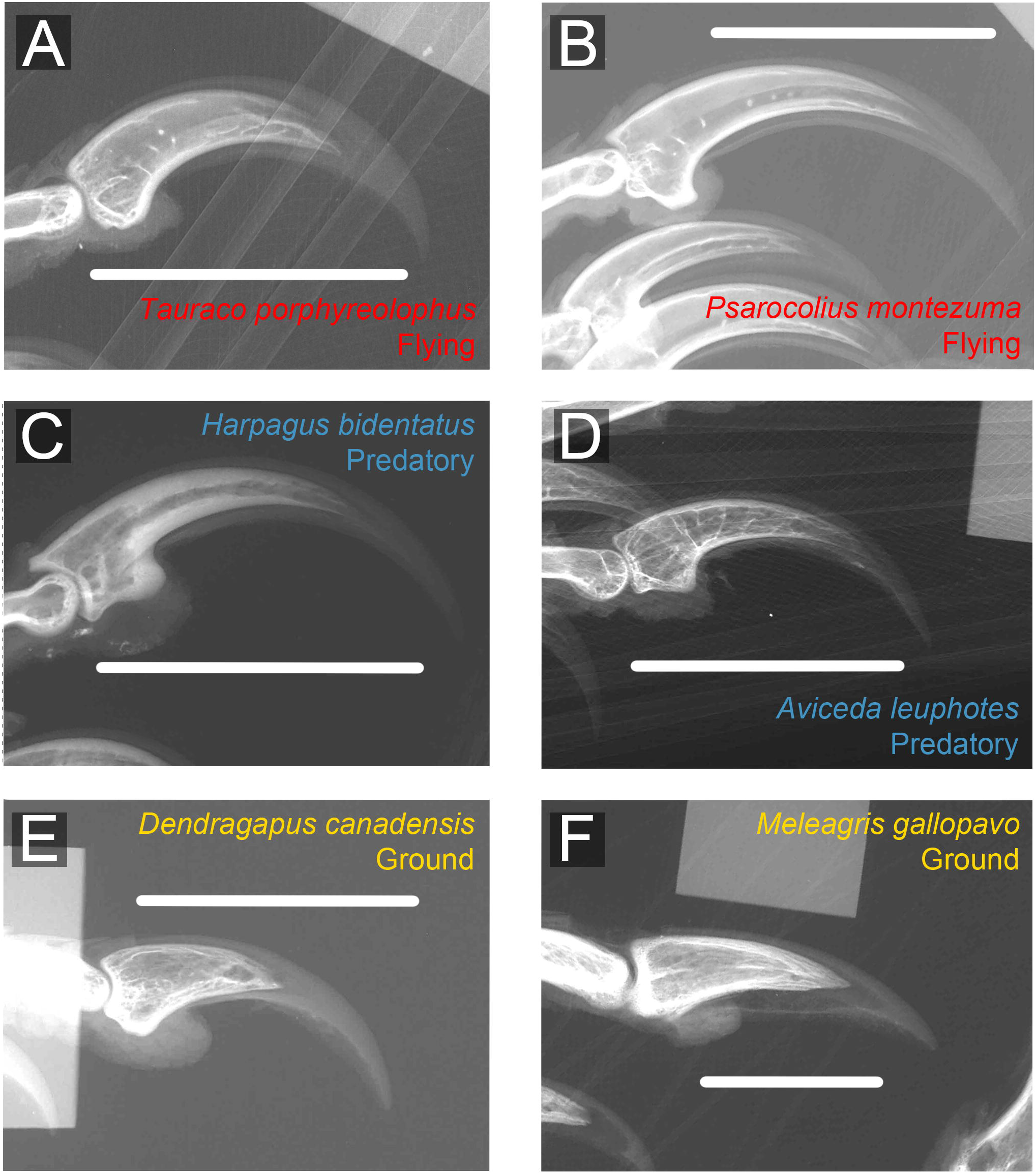
Representative third pedal unguals showing typical claws for each of the three ecological groups. Flying taxa: (A) *Tauraco porphyreolophus* (Purple-crested Tauraco) and (B) *Psarocolius montezuma* (Black Oropendola); Predatory taxa: (C) *Harpagus bidentatus* (Double-toothed Kite) and (D) *Aviceda leuphotes* (Pacific Baza); Cursorial taxa: (E) *Dendragapus canadensis* (Blue Grouse) and (F) *Meleagris gallopavo* (Wild Turkey).

### 2.1 Ecology and Shape

Three traditional morphometric measures were taken from claw radiographs: (1) the ratio of length of the bony core to length of keratinous sheath; (2) the dorsal arc of the bony core (Figure 2A); and (3) the dorsal arc of the keratinous sheath (Figure 2B). These measurements were taken from radiographs in ImageJ (Schneider et al., 2012) following the general scheme set forth by Pike and Maitland (2004). The ventral arc of the claw was not calculated, given that many claws have a ventral constriction near the claw tip (Pike & Maitland, 2004). Note that these traditional methods assume that both the arcs of the bony core and keratinous sheath inscribe a circle. Seven points were measured on each claw: the tip of the keratinous sheath, the tip of the bony core, the midpoint of the crescent-shaped articulation surface with the penultimate phalanx, the dorsal lip of the bony core, the dorsal lip of the keratinous sheath, and approximate midpoints along the arcs of the bony core and keratinous sheath. These points were used to calculate the center of the circle that the bony core and keratinous sheaths inscribe, which were then used to calculate arcs following previous studies (Feduccia, 1993; Pike & Maitland, 2004).

**FIGURE 2.**
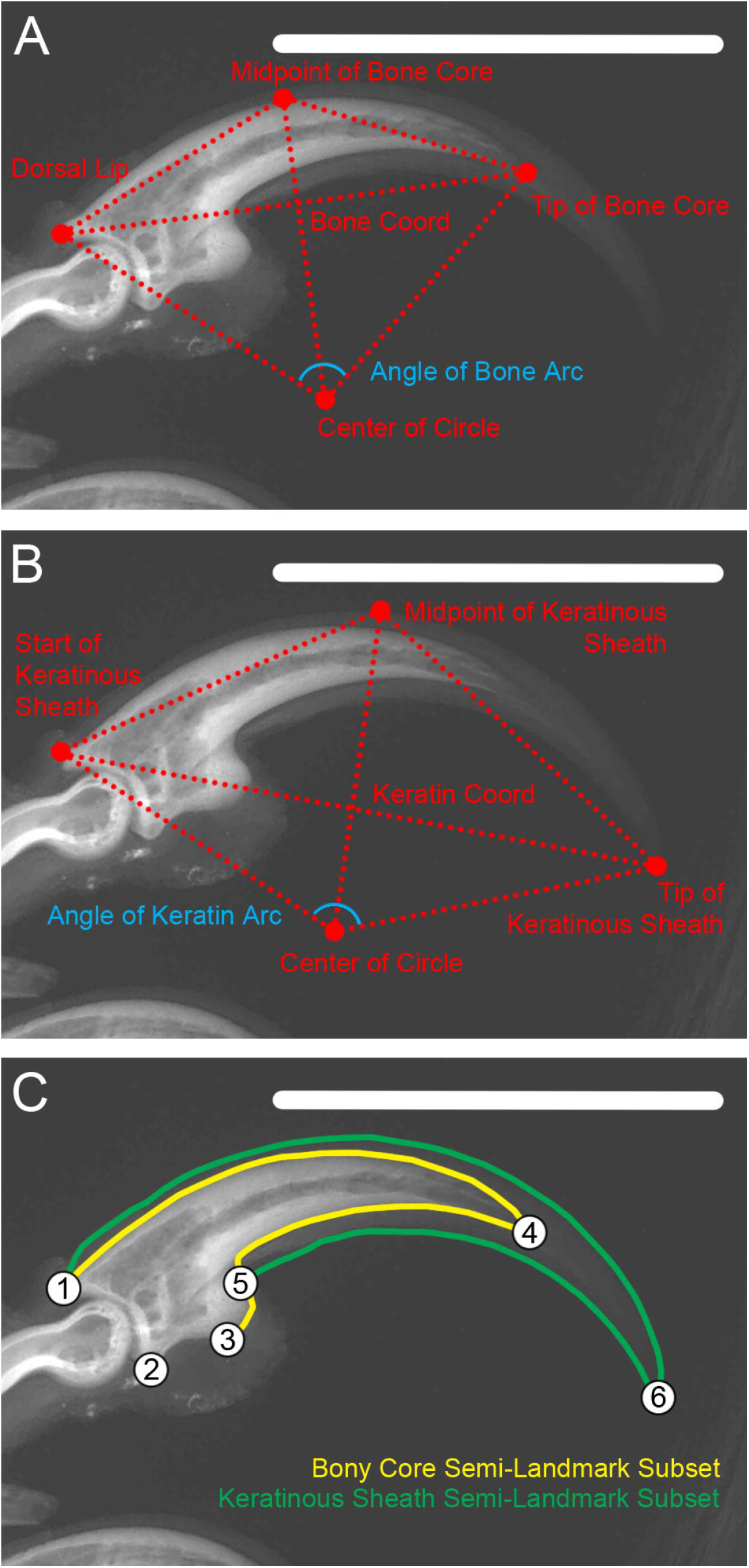
Traditional arc measurements taken for the (A) bony core and (B) keratinous sheath. (C) Landmark configuration with numbered landmarks and semi-landmark curves for the bony core (yellow) and keratinous sheath (green). Landmark definitions in supplement.

For geometric morphometric analyses, six landmarks and 77 semi-landmarks were digitized onto the radiographs (Figure 2C) in the tpsDig2 software (Rohlf, 2006). The goal underlying landmark selection was to capture both the shape of the bony core and the keratinous sheath. Landmark numbers and definitions are listed in the supplement (Table S1). The landmark data were imported into the R package *geomorph* (Adams & Otárola-Castillo, 2013) and subjected to Generalized Procrustes Analysis (GPA). The bending energy criterion was used when sliding semi-landmarks (Perez et al., 2006).

The amount of variance and range of values for each traditional metric was calculated for the entire dataset and for four species which had the largest sample sizes among the data to assess individual variation surrounding species means (*Milvus migrans*, Accipitriformes, n = 19; *Dendragapus canadensis*, Galliformes, n = 20; *Puffinus griseus*, Procellariformes, n = 21; *Tinamus major*, Tinamiformes, n = 25). For geometric morphometric data, a principal component analysis (PCA) was run on all data (n = 580) highlighting these four species to visualize the impact of individual variation on morphospace occupation.

To assess the relationship between traditional claw metrics and ecological groups quantitatively, phylogenetic generalized least squares (PGLS) regressions using maximum likelihood estimates of Pagel’s lambda (Pagel, 1999) were run in the *nlme* package in R (Pinheiro et al., 2018; R Core Team, 2018). This serves to include the amount of relevant phylogenetic information in the model and does not assume Brownian motion or a star phylogeny. Pairwise comparisons were performed on species means for each traditional metric using *phytools* in R (Revell, 2012). All PGLS models included size as a factor, as some previous studies have found a significant impact of size on claw arc metrics (Pike & Maitland, 2004; Birn-Jeffery et al., 2012; Csermely et al., 2012). While body mass is commonly used as a size metric, museum specimens do not usually have body mass recorded and only 89% of the included taxa had known body masses in the literature (Dunning et al., 2007). Further, sex was unknown for some of the specimens included in this study and sexual size dimorphism is large in many of the sampled birds (e.g., *Meleagris gallopavo*). Therefore, taking the average of male and female body masses would likely have led to poor estimates of actual body mass for the specimens. As an alternative, claw centroid size –the square root of the sum of squared inter-landmark distances– was used as the body mass metric. Previous studies have found that the size of pedal digit III claws and body masses are tightly correlated in birds regardless of ecological mode over a wide range of body masses (Pike & Maitland, 2004).

To evaluate geometric morphometric data, a PCA was run on species means to distinguish between taxon trends in morphospace. The impacts of phylogeny were assessed using the multivariate version of Blomberg’s K statistic (Blomberg et al., 2003; Adams, 2014). The degree of allometric signal in the data was determined by testing for a correlation between the common allometric component and log-transformed centroid size of the claws (Mitteroecker et al., 2004). Given a significant allometric signal, a phylogenetic Procrustes ANOVA (Goodall, 1991) was run testing the relationship between shape, size, and ecological group. Finally, differences in levels of disparity were evaluated using Procrustes variance as the disparity metric (Zelditch et al., 2012) using 999 permutations to calculate significance.

### 2.2 Relationship between the bony core and keratinous sheath

PGLS models were performed on species means with log-transformed centroid size as a covariate to test for significant correlations between the log-transformed bony core and log-transformed keratinous sheath arcs. The R^2^ coefficient was used to assess the amount of variance of the bony core arc that explained the keratinous sheath arc. Then, a phylogenetic ANOVA with pairwise comparisons of the residuals from the above PGLS and ecological group was used to determine whether this relationship was different among the three ecological groups. Finally, a phylogenetic paired t-test was run comparing the bony core arc measurements with the keratinous sheath arc measurements to test if the bony core arcs and keratinous sheath arcs were statistically different from one another.

For geometric morphometric data, claw landmarks were placed into two separate subsets following GPA. Landmarks 1, 2, 3, 4, and 7–37 were assigned to the bony core subset and landmarks 5, 6, and 38–83 were assigned to the keratinous sheath subset (Figure 2C). Hypotheses of modularity and integration were then tested for species means accounting for the impacts of phylogeny under a Brownian motion model of evolution in *geomorph* (Adams & Felice, 2014; Adams, 2016; Adams & Collyer, 2016). The covariance ratio (CR) coefficient was calculated from the data and then compared to a null distribution of CR values based on landmarks being randomly assigned to the two landmark subsets for 999 iterations. When the observed CR coefficient is significantly lower than the null distribution, the hypothesis of modularity is supported. Integration was evaluated using phylogenetic partial least squares (PLS) analysis. Significance was determined by randomly permuting landmarks in the two landmark subsets for 999 iterations. Differences in integration between ecological groups were then compared using the compare.pls function in *geomorph*.

### 2.3 Comparison of traditional and geometric morphometric methods

Finally, two-block PLS analysis was employed to assess the degree of similarity between the traditional and geometric morphometric data. The three traditional morphometric measures (ratio of bony core length to keratinous sheath length, log-transformed bony core arc, log-transformed keratinous sheath arc) were combined into a single block and the geometric morphometric shape data were combined into the second block. As above, phylogenetic two-block PLS analyses was run using species means and 999 iterations in *geomorph*.

## 3 Results

### 3.1 Ecology and Shape

All three traditional morphometric measures had relatively low variance across all taxa. The bony core-keratinous sheath length ratio confidence intervals were from 0.693–0.705, the bony core arc ranged from 74.6°–77.9°, and the keratinous sheath arc ranged from 102.2°–107° (Figure 3A). When assessing intraspecific variance, the four highly sampled taxa had confidence intervals that were a maximum of 2–4.1% wide for the length ratio, 2.64–7.08° wide for the bony core arc, and 4.66-10.3° wide for the keratinous sheath arc (Figure 3C–F; Table S4). While these confidence intervals were not large in absolute terms, they suggest that individuals of the same species may be more different from one another than individuals of different, related species. The geometric morphometric data demonstrate that each of these four taxa cluster intraspecifically in PCA, but that each taxon ranges across morphospace such that the intraspecific variation is often larger than interspecific variation (Figure 3B; Table S2).

**FIGURE 3.**
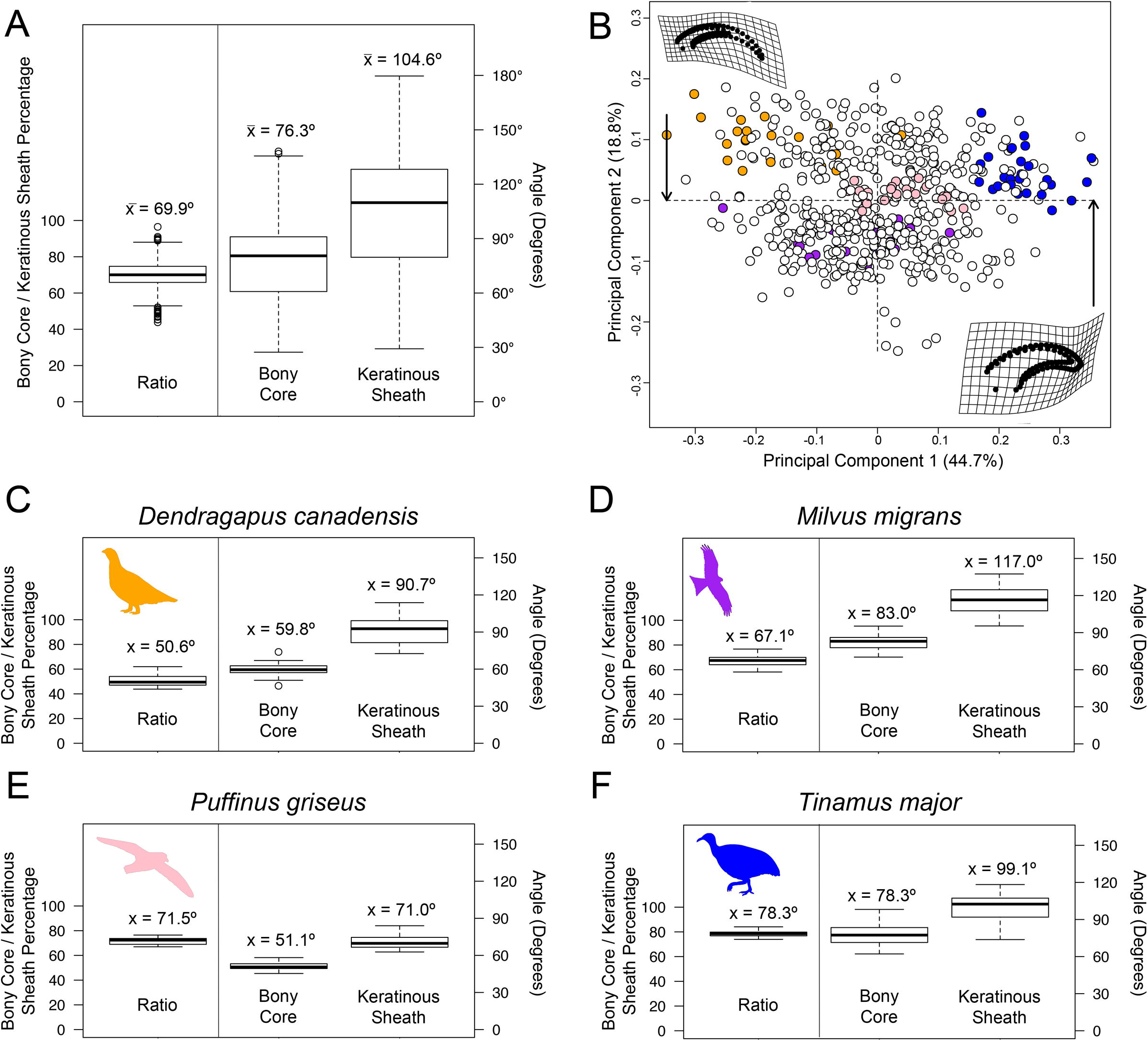
Range of variation for (A) traditional measurements and (B) geometric morphometric data. Orange - *Dendragapus canadensis* (n = 20); Purple – *Milvus migrans* (n = 19); Pink – *Puffinus griseus* (n = 21); Dark blue - *Tinamus major* (n = 25). Variation in traditional morphometric metrics for (C) *Dendragapus canadensis*, (D) *Milvus migrans*, (E) *Puffinus griseus*, and (F) *Tinamus major*. Both traditional and geometric morphometric data demonstrate substantial intraspecific variation in claw shape.

Phylogenetic ANOVAs revealed that ecological groups are not significantly correlated with the length ratio (p = 0.272), log-transformed bony core arc (p = 0.489), or log-transformed keratinous sheath arc (p = 0.314) after accounting for body size. Similar to the traditional metrics, geometric morphometric data did not show a significant difference in claw shape across groups in spite of apparent groupings in PCA (Table S5).

For geometric morphometric means data, principal component 1 (PC1) summarized 47.7% of total variance and PC2 summarized 18.9% of total variance (Table S3). Although PC1 revealed separation between ground-dwelling and predatory taxa (Figure 4A), flying generalists ranged broadly across morphospace. Taxa on the positive end of PC1 had blunt short claws in which the bony core to keratinous sheath length ratio was high, similar to ground-dwelling taxa (Figure 4B). Taxa on the negative end of PC1 had claws with a strong recurve similar to predatory taxa with a bony core to keratinous sheath length ratio closer to 0.70. PC2 did not separate the three ecological groups. The consensus shape for the positive end of the PC2 axis had a slight recurve and a high bony core to keratinous sheath ratio. The negative end of PC2 was characterized by a flattened, elongate claw. The geometric morphometric data were significantly correlated with phylogeny (K_mult_ = 0.155, p < 0.001) and allometry (Figure 4C), albeit with a low percent of shape variance explained by allometry (R^2^ = 0.03). Procrustes variance was roughly the same for all three groups (flying generalists = 0.71, ground dwellers = 0.66, predatory = 0.72), with no groups having significantly different levels of disparity.

**FIGURE 4.**
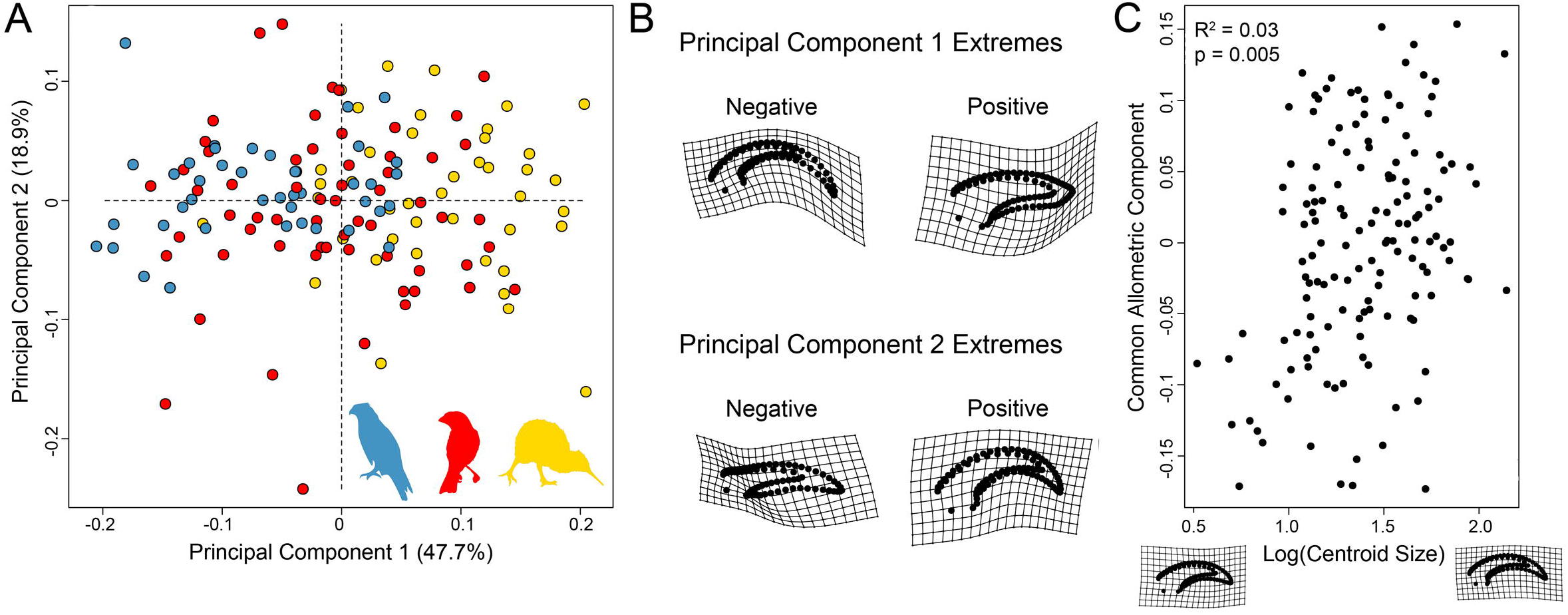
Geometric morphometric claw shape data. (A) Principal component analysis of total claw shape showing separation between predatory and ground birds with flying birds spreading across morphospace. Blue = predatory, red = flying, yellow = ground. (B) Thin-plate spline (TPS) representations of the positive and negative extremes of PC1 and PC2. (C) Allometric analysis of the common allometric component of shape and log-transformed centroid size. TPS grids show representations of small (left) and large (right) claw shape.

### 3.2 Relationship between the bony core and keratinous sheath

A PGLS of log-transformed keratinous sheath arc and log-transformed bony core arc data including body size as a factor had a significant association (λ = 0.702, m = 0.873, p < 0.001) with a substantial amount of total variance of the bony core arc explaining the keratinous sheath arc (R^2^ = 0.792) (Figure 5A). Residuals of this PGLS and ecological group did not reveal a significant association (p = 0.185) suggesting that different ecologies did not have differing trajectories between their bony core arc and keratinous sheath arc. A t-test did show that the log-transformed bony core arc and log-transformed keratinous sheath arc are significantly different from one another (p < 0.001) with the keratinous sheath having a larger angle than the bony core.

**FIGURE 5.**
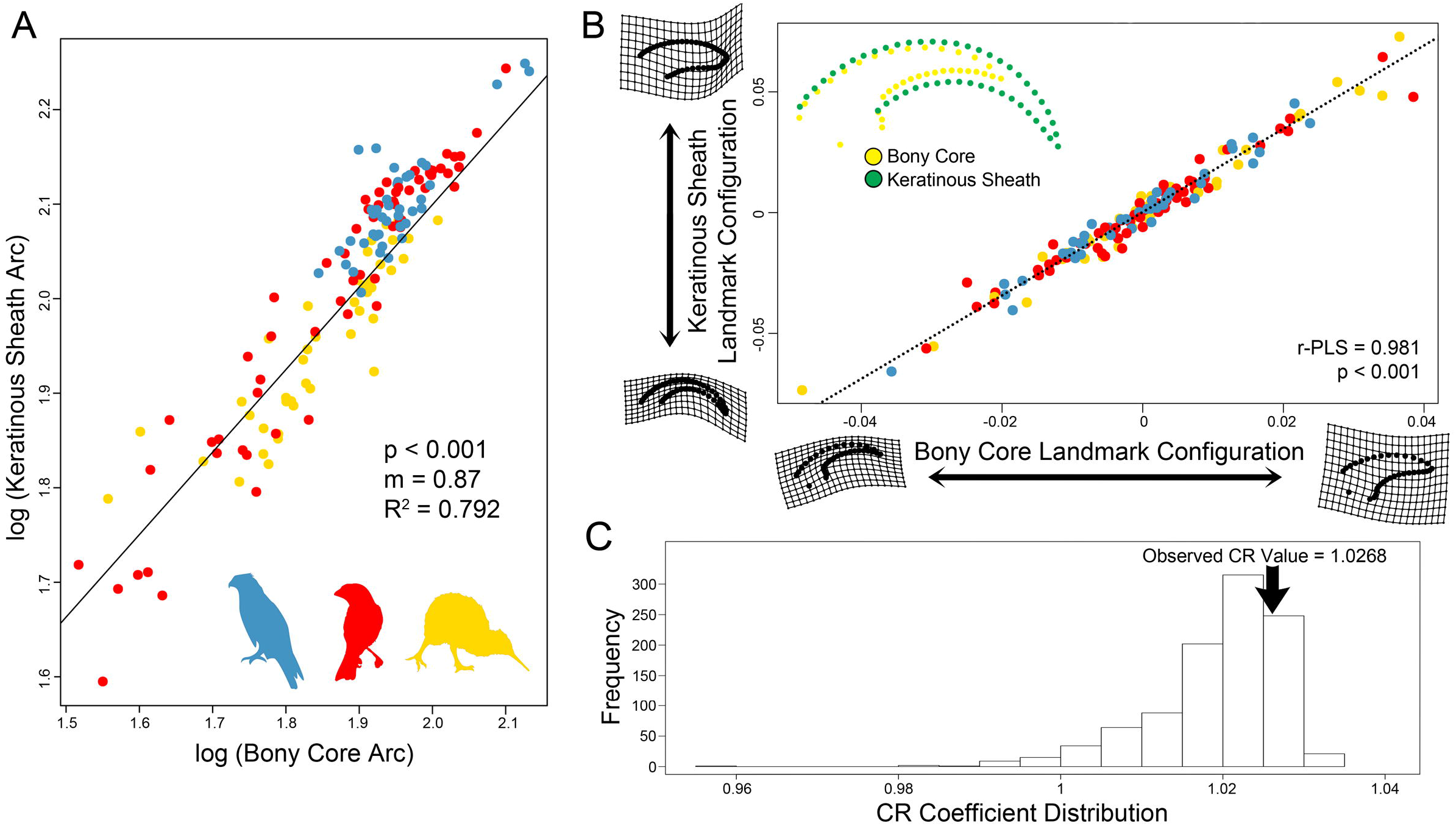
Measures of integration between the bony core and keratinous sheath for both (A) traditional morphometric data using phylogenetic general least squares regression (blue = predatory, red = flying, yellow = ground) and (B) geometric morphometric data using two-block partial least squares analysis. TPS grids show differences in shape along each shape block. (C) Assessment of modularity showing the observed covariance ratio (CR) against a null distribution. The observed CR is not significantly lower than the distribution and so modularity is not supported.

Using geometric morphometric data, the phylogenetically informed integration analysis supported significant integration between the bony core and keratinous sheath (r-PLS = 0.981, p < 0.001). Taxa with recurved bony cores had recurved keratinous sheaths and taxa with flattened, short bony cores had flattened, short keratinous sheaths (Figure 5B). Further, the phylogenetically informed analysis of modularity did not support the bony core and keratinous sheath as separate modules (CR = 1.02, p = 0.845) (Figure 5C). Comparison of integration levels did not suggest that any ecological group was significantly more integrated than any other group (Table S5).

### 3.3 Comparison of traditional and geometric morphometric methods

Finally, a phylogenetic PLS of traditional metrics and geometric morphometric shape data had a significant, but loose correlation (r-PLS = 0.506, p < 0.001, Figure 6). This suggests that the two types of data summarize claw shape in somewhat complementary ways, but that the data types are not strongly integrated.

**FIGURE 6.**
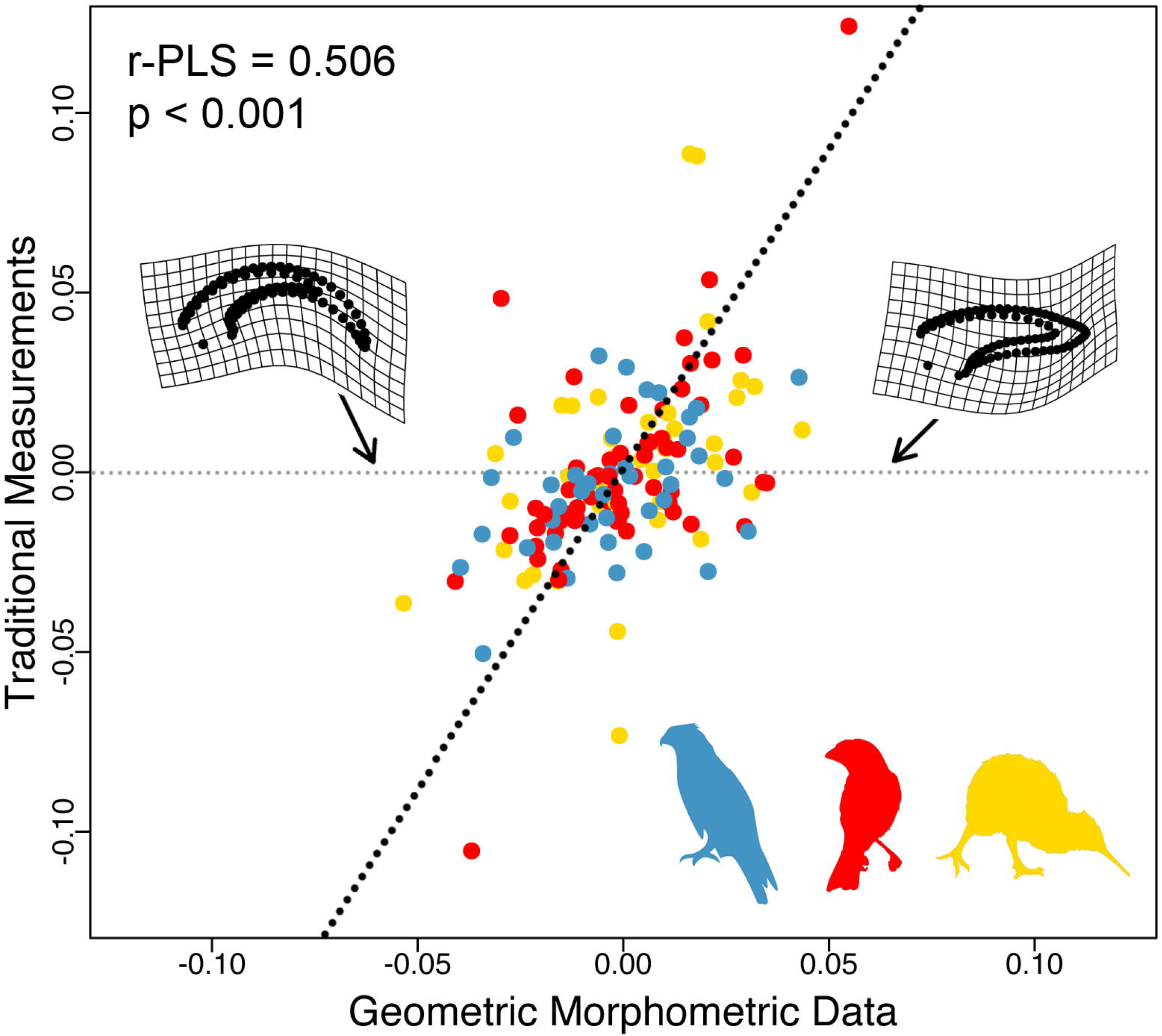
Two-block partial least squares analysis of geometric morphometric data against traditional morphometric data. Inset TPS grids show shapes at the positive and negative ends of the geometric morphometric block. Blue = predatory, red = flying, yellow = ground.

## 4 Discussion

Claw arc measurements were first used in the early 1990s, with the goal of predicting the ecology of fossil bird taxa based on the relationship between claw shape and ecology in extant birds (Peters and Görgner, 1992; Feduccia, 1993). Since that pioneering work, some studies have found a strong correlation between bird claw shape and ecology (Glen & Bennett, 2007; Csermely et al., 2012; Cobb & Sellers, in review) while others have not (Pike & Maitland, 2004; Birn-Jeffery et al., 2012). These conflicting results are exacerbated by the lack of overlap in taxa and methodologies across studies (Birn-Jeffrey et al., 2012). Additionally, many paleobiologists have used keratinous sheath shape in extant taxa to attempt to reconstruct ecological mode in extinct taxa using bony core shape, which have thus far been assumed to be complementary. After incorporating phylogeny and body size, we found that neither traditional nor geometric morphometrics recovered significant differences in claw shape between ecological categories. The bony core and keratinous sheath are significantly integrated and the degree of integration does not differ across ecological groups, but the bony core shape and keratinous sheath shape are significantly different and cannot be compared without corrections.

Although body mass was found to be significantly correlated with claw shape, we support previous studies in asserting that body size is not a substantial predictor of claw shape (Pike & Maitland, 2004; Birn-Jeffrey et al., 2012; Cobb & Sellers, in review) given that only 3% of total variance is explained by size (Figure 4B). The relationship between body mass and claw arc has been shown to be complex, wherein the relationship likely varies within ecological categories (Pike & Maitland, 2004; Birn-Jeffrey et al., 2012). For example, claw angle increases with body mass in predatory and climbing birds, but decreases with body mass in ground birds, and is not correlated with body mass in perching birds (Pike & Maitland, 2004). Therefore, it is important to include a size correlate in models even if the overall variation explained by size is small. Studies that do not include size as a predictor variable likely introduce confounding effects.

Similar to body size, phylogenetic relatedness is a likely confounding effect in any comparative study (Felsenstein, 1985) and differing relationships between claw shape and ecology in previous studies may be due to a lack of the consistent application of phylogenetic comparative methods. A strong phylogenetic signal was uncovered for claw shape whether it was derived using traditional or geometric morphometrics, demonstrating the importance of using phylogenetic comparative methods when examining claw shape. While some recent studies (Cobb & Sellers, in review) have argued that phylogenetic comparative methods cannot be used for comparative studies of birds, citing recent conflicting bird trees, birds are one of the best vertebrate groups for the application of phylogenetic comparative methods, with many complete trees that largely agree (Jetz et al., 2012; Prum et al., 2015). Previous claw studies have found that morphological trends follow family level groupings in the absence of phylogenetic comparative methods (Fowler et al., 2009) and that the implementation of independent contrasts largely eliminates the significant relationship between claw geometry and behavior (Birn-Jeffery et al., 2012). Phylogenetic comparative methods must be employed when assessing the relationship between claw shape and ecology in birds due to substantial ecological convergence in groups separated by long branch lengths.

Although there were no significant associations between ecology and shape, the PCA of geometric morphometric shape data did show some separation between ecological groups, especially between predatory and ground birds. The generalist flying category was spread widely across morphospace, invading the predatory and ground bird regions of morphospace (Figure 4A). It is therefore possible that our groupings were too broad to detect ecological differences, especially because members of our groups may have used their claws in non-complementary ways. For example, both Strigiformes and Accipitriformes were placed in the predatory category, but previous studies have found them to strongly separate in claw shape morphospace due to different prey capture techniques (Csermely et al., 2012). Still, there were not significant differences between groups that were very ecologically distinct (ground and predatory birds) after incorporating phylogeny and body size even though they were clearly separated in morphospace (Figure 4A). Based on morphospace occupation, it is possible that geometric morphometrics may discriminate claw shape across ecological groups better than traditional morphometrics, especially considering that these two data types are only loosely correlated with one another (Figure 6).

Additionally, there was a previously unrecognized confounding factor present in the data, high intraspecific variation. Some previous studies used up to six specimens per species when examining claw shape (Birn-Jeffery et al., 2012) while some other studies have used a single individual per species (Csermely et al., 2012). When examining several species that had high sample sizes in the dataset (n > 18), very high intraspecific variation is documented, whether using traditional measurements or geometric morphometric analyses (Figure 3). Plotting four species with large sample sizes in morphospace showed some within-species clustering, but clusters ranged quite widely, often taking up a large portion of occupied morphospace (Figure 3B). Ethier et al. (2010) warned workers not to use a single bird species in studies of claw development or morphology as a result of variable rates of claw growth due to fluctuating energy demands such as migration and breeding. Cobb and Sellers (in review) found that the left and right claws of the same fossil taxon could be categorized in different ecological categories. Although this may also be related to taphonomy (Hedrick & Dodson, 2013; Hedrick et al., 2019), no study has thus far looked at within-individual variation in claw shape in extant or fossil birds. It is likely that intraspecific variation is an additional, previously unappreciated factor that may have led to discrepancies in results in prior studies.

The size and shape of the bony core and keratinous sheath of avian claws has previously been considered to be similar (Clark, 1936; Ethier et al., 2010), but this relationship had not been tested. The bony core and keratinous sheath were significantly and strongly correlated, suggesting they act as a functional unit, using both traditional morphometric (Figure 5A) and geometric morphometric approaches (Figures 5B, 5C). This is important to establish based on the large amount of work evaluating ecology in extinct taxa by comparing the bony cores of extinct taxa to the keratinous sheaths of extant taxa. Glen and Bennett (2007) presented the first estimation of the keratinous sheath arc from the bony core arc using radiographs, but they did not present the strength of the correlation, only the conversion factor. Recently Cobb and Sellers (in review) used radiographs to determine whether the bony ungual or keratinous sheath was a better predictor of avian ecology, but they did not test for the degree of integration between the two data types. We found that the log-transformed bony core arc explains 79.2% of the variation in the log-transformed keratinous sheath arc when incorporating phylogeny and body size, suggesting a relatively tight correlation (Figure 5A). The geometric morphometric data had even stronger levels of integration between the bony core and keratinous sheath (Figures 5B; 5C). These results fit with similar studies exploring covariation between the keratinous rhamphotheca and underlying bone of bird beaks that document shape covariation both when considering (Button & Zanno. in review) and disregarding (Urano et al. 2018) phylogenetic relatedness. These data suggest that predicting keratinous sheath shape from bony core shape is feasible and is better done using geometric morphometrics than traditional morphometrics. However, the bony core shape and keratinous sheath shape are statistically different from one another in spite of being correlated and therefore it is inappropriate to directly compare the bony core of fossil claws with the keratinous sheath of extant taxa without a conversion factor.

As first noted by Pike and Maitland (2004), one limitation of this study and all previous studies is that these analyses were all run on two-dimensional representations of three-dimensional structures. Predatory birds often have conical, tapering claws whereas climbing birds have laterally compressed claws with sharp distal ends (Richardson, 1942; Yalden, 1985; Peters & Görgner, 1992; Pike & Maitland, 2004). Throughout the course of this work, we noticed numerous claws that were mediolaterally asymmetric, such as those of some woodpecker and parrot species. Although this has yet to be reported in a quantitative framework, it is likely related to function. This information is completely lost when quantifying shape using photographs or radiographs and future claw shape studies using CT will be necessary to better categorize birds into distinct ecological categories.

## 5 Conclusion

After accounting for body size and phylogenetic relatedness, neither traditional morphometrics nor geometric morphometrics are capable of significantly separating birds into ecological categories. However, ordination analyses do demonstrate some separation in morphospace when using geometric morphometrics, suggesting that geometric morphometrics improves on traditional morphometric methods at ecological classification (Schmieder et al., 2015; Hedrick et al., 2018). Although there has been a large amount of previous work on claw shape, differing methods of data collection and analysis have precluded a consensus on how avian ecology impacts claw shape. We advocate for geometric morphometrics of claw radiographs or CT scans as a promising new method for characterizing claw shape and strongly suggest that future studies incorporate confounding factors such as body size, phylogeny, and individual variation before assessing ecology in extinct taxa.

## ACKNOWLEDGEMENTS

We thank Nathan Rice and Ned Gilmore (both Academy of Natural Sciences of Drexel University–ANSP) for collections access and John Lundberg and Kyle Luckenbill (both ANSP) for access to and instruction in the use of the ANSP x-ray facilities. We thank Wilfried Mai for access to the x-ray facilities at the Ryan Veterinary Hospital of the University of Pennsylvania for a proof of concept study. Jane Dmochowski (UPenn) gave valuable discussions on earlier versions of this manuscript. Finally, we thank the Greg and Susan Walker Endowment (awarded to S. A. C.), the University of Pennsylvania Paleobiology Summer Stipend (S. A. C.), and the National Science Foundation (NSF 1612211–awarded to B. P. H.) for funding.

## CONFLICT OF INTEREST

The authors declare no conflict of interest.

## AUTHOR CONTRIBUTIONS

BPH, SAC, LEZ, CN, and PD conceived of the study. SAC collected the data and landmarked the claws. BPH and SAC analyzed the data. BPH, SAC, LEZ, CN, and PD wrote the manuscript.

## DATA ACCESSIBILITY

Upon article acceptance, original data including radiographs and supplemental information will be archived into the public repository Dryad.

---

**Figure S1.** Tree based on Jetz et al. (2012) showing the phylogenetic relationships between the 145 taxa used in the study. Tips are colored by locomotor mode showing numerous convergences in mode across taxa studied. Blue = predatory, red = flying, yellow = ground.

**Supplemental Table 1:** Landmarks, semi-landmarks, and landmark definitions for geometric morphometric data. Bookstein’s topology of landmarks are noted for each landmark. Further, each landmark is divided into either the bony core or keratinous sheath subset for integration and modularity analyses.

**Supplemental Table 2:** Specimen number, species name, family, traditional morphometric metrics, and the first 10 PC scores for geometric morphometric data for all 580 individuals.

**Supplemental Table 3:** Specimen number, species name, family, centroid size mean, traditional morphometric metrics, and the first 10 PC scores for geometric morphometric data for species means (n = 145).

**Supplemental Table 4:** Statistical results for traditional morphometric analyses. Mean, standard deviation, and confidence intervals for intraspecific analyses. PGLS of log-transformed keratinous sheath arc and log-transformed bony core arc. Phylogenetic ANOVAs for traditional morphometrics. T-test comparing bony core and keratinous sheath log-transformed arc measurements.

**Supplemental Table 5:** Statistical results for geometric morphometric analyses. Tests for allometry and phylogenetic signal. Claw shape by group including centroid size as a covariate and pairwise comparisons among ecological groups. Disparity, integration, and modularity analyses.

## REFERENCES

Adams, D.C. (2014). Quantifying and comparing phylogenetic evolutionary rates for shape and other high-dimensional phenotypic data. Systematic Biology 63, 166–177.

Adams, D.C. (2016). Evaluating modularity in morphometric data: Challenges with the RV coefficient and a new test measure. Methods in Ecology and Evolution 7, 565–572.

Adams, D.C. & Felice, R.N. (2014). Assessing trait covariation and morphological integration on phylogenies using evolutionary covariance matrices. PLoS One, e94335.

Adams, D.C. & Otárola-Castillo, E. (2013). geomorph: an R package for the collection and analysis of geometric morphometric shape data. Methods in Ecology and Evolution 4, 393–399.

Adams, D.C. & Collyer, M.L. (2016). On the comparison of the strength of morphological integration across morphometric datasets. Evolution 70, 2623–2631.

Birn-Jeffery, A.V., Miller, C.E., Naish, D., Rayfield, E.J. & Hone, D.W.E. (2012). Pedal Claw Curvature in Birds, Lizards and Mesozoic Dinosaurs – Complicated Categories and Compensating for Mass-Specific and Phylogenetic Control. PLoS ONE 7, e50555.

Blomberg, S.P., Garland, T. & Ives, A.R. (2003). Testing for phylogenetic signal in comparative data: behavioral traits are more labile. Evolution 57, 717–745.

Bookstein, F.L. (1991). Morphometric tools for landmark data: geometry and biology. New York: Cambridge University Press.

Button K. & Zanno, L.E. (in review) Bone shape and surficial texture predict rhamphotheca morphology in birds.

Clark, W.E.L. (1936) The problem of the claw in primates. Proceedings of the Zoological Society of London 1, 1–24.

Cobb, S.E. & Sellers, W.I. (in review). Inferring lifestyle for Aves and Theropoda: a model based on curvatures of extant avian ungual bones. bioRxiv, https://doi.org/10.1101/517375.

Corti, M. (1993). Geometric morphometrics: an extension of the revolution. TREE 8, 302–303.

Csermely, D., Rossi, I. & Nasi, F. (2012). Comparison of claw geometrical characteristics among birds of prey and non-raptorial birds. Italian Journal of Zoology 79, 410–433.

Csermely, D. & Rossi, O. (2006). Bird claws and bird of prey talons: Where is the difference? Italian Journal of Zoology 73, 43–53.

Dunning Jr., J.B. (1993). Body masses of birds of the world. Ann Arbor, MI, USA: CRC Press.

Dunning Jr., J.B. (2007). CRC Handbook of Avian Body Masses, second edition. Boca Raton, Florida: CRC Press.

Ethier, D.M., Kyle, C.J., Kyser, T.K. & Nocera, J.J. (2010). Variability in the growth patterns of the cornified claw sheath among vertebrates: implications for using biogeochemistry to study animal movement. Canadian Journal of Zoology 88, 1043–1051.

Feduccia, A. (1993). Evidence from Claw Geometry Indicating Arboreal Habits of Archaeopteryx. Nature 259, 790–793.

Felsenstein, J. (1985). Phylogenies and the comparative method. The American Naturalist 125, 1–15.

Fowler, D.W., Freedman, E.A. & Scannella, J.B. (2009). Predatory Functional Morphology in Raptors: Interdigital Variation in Talon Size Is Related to Prey Restraint and Immobilisation Technique. PLoS ONE 4, e7999.

Fowler, D.W., Freedman, E.A., Scannella, J.B. & Kambic, R.E. (2011). The Predatory Ecology of Deinonychus and the Origin of Flapping in Birds. PLoS ONE 6(12), e28964.

Glen, C.L. & Bennett, M.B. (2007). Foraging modes of Mesozoic birds and non-avian theropods. Current Biology 17, R911–R912.

Goodall, C.R. (1991). Procrustes methods in the statistical analysis of shape. J. R. Stat. Soc. B. 53, 285–339.

Hedrick, B.P. & Dodson, P. (2013). Lujiatun psittacosaurids: Understanding individual and taphonomic variation using 3D geometric morphometrics. PLoS ONE 8, e69265.

Hedrick, B.P. & Dumont, E.R. (2018). Putting the leaf-nosed bats in context: A geometric morphometric analysis of three of the largest families of bats. Journal of Mammalogy 99(5), 1042–1054.

Hedrick, B.P., Schachner, E.R., Rivera, G., Dodson, P. & Pierce, S.E. (2019). The effects of skeletal asymmetry on interpreting biologic variation and taphonomy in the fossil record. Paleobiology 45(1), 154–166.

Jetz, W., Thomas, G.H., Joy, J.B., Hartmann, K. & Mooers, A.O. (2012). The global diversity of birds in space and time. Nature 491, 444–448.

Mitteroecker, P., Gunz, P., Bernhard, M., Schaefer, K. & Bookstein, F.L. (2004). Comparison of cranial ontogenetic trajectories among great apes and humans. Journal of Human Evolution 46, 679–698.

Mitteroecker, P. & Gunz, P. (2009). Advances in geometric morphometrics. Evolutionary Biology 36, 235–247.

Pagel, M. (1999). Inferring the historical patterns of biological evolution. Nature 401, 877–884.

Perez, S.I., Bernal, V. & Gonzalez, P.N. (2006). Differences between sliding semi-landmark methods in geometric morphometrics, with an application to human craniofacial and dental variation. Journal of Anatomy 208, 769–784.

Peters, S.F. & Görgner, E. (1992). A comparative study on the claws of Archaeopteryx. In Papers in Avian Palaeontology: 29–37. Campbell, K. (Ed.). Los Angeles, CA: Natural History Museum of Los Angeles County.

Pike, A.V.L. & Maitland, D.P. (2004). Scaling of Bird Claws. Journal of Zoology, London 262, 73–81.

Pinheiro, J., Bates, D., DebRoy, S., Sarkar, D. & R Core Team. (2018). nlme: Linear and Nonlinear Mixed Effects Models. R Package Version.

Prum, R.O., Berv, J.S., Dornburg, A., Field, D.J., Townsend, J.P., Lemmon, E.M. & Lemmon, A.R. (2015). A comprehensive phylogeny of birds (Aves) using targeted next-generation DNA sequencing. Nature 526, 569.

R Core Team. (2018). R: A language and environment for statistical computing. R Foundation for Statistical Computing, Vienna, Austria. URL https://www.R-project.org/.

Revell, L.J. (2012). An R package for phylogenetic comparative biology (and other things). Methods of Ecology and Evolution 3, 217–223.

Richardson, F. (1942). Adaptive modifications for tree-trunk foraging in birds. University of California Publications in Zoology 46, 317–367.

Rohlf, F.J. (2006). tpsDig, digitize landmarks and outlines, version 2.05. Department of Ecology and Evolution, State University of New York, Stony Brook, New York.

Schmieder, D.A., Benítez, H.A., Borissov, I.M. & Fruciano, C. (2015). Bat species comparisons based on external morphology: a test of traditional versus geometric morphometric approaches. PLoS ONE 10, e0127043.

Schneider, C.A., Rasband, W.S. & Eliceiri, K.W. (2012). NIH Image to ImageJ: 25 years of image analysis. Nature methods 9(7), 671–675.

Slice, D.E. (2007). Geometric morphometrics. Annual Review of Anthropology 36, 261–281.

Urano, Y., Tanoue, K., Matsumoto, R., et al. (2018) How does the curvature of the upper beak bone reflect the overlying rhinotheca morphology? Journal of Morphology 28, 788. DOI: 10.1002/jmor.20799.

Yalden, D.W. (1985). Forelimb function in *Archaeopteryx*. In The Beginnings of Birds: 91–97. Hecht, M.K., Ostrom, J., Viohl, G. & Wellnoster, P. (Eds.). Eichstatt, Germany: Freunde des Jura-Museums.

Zani, P.A. (2000). The comparative evolution of lizard claw and toe morphology and clinging performance. Journal of Evolutionary Biology 13, 316–325.

Zelditch, M.L., Swiderski, D.L., Sheets, H.D. & Fink, W.L. (2012). Geometric Morphometrics for Biologists: A Primer. 2 edn. London, UK: Elsevier Academic Press.

